# RNA X-ray footprinting reveals the consequences of an *in vivo* acquired determinant of viral infectivity

**DOI:** 10.1101/2021.08.10.455819

**Authors:** Rebecca Chandler-Bostock, Richard J. Bingham, Sam Clark, Andrew J. P. Scott, Emma Wroblewski, Amy Barker, Simon J. White, Eric C. Dykeman, Carlos P. Mata, Jen Bohon, Erik Farquhar, Reidun Twarock, Peter G. Stockley

## Abstract

The secondary structures of the bacteriophage MS2 ssRNA genome, frozen in defined states, were determined with minimal perturbation using constraints from X-ray synchrotron footprinting (XRF). The footprints of the gRNA in the virion and as transcript are consistent with single, dominant but distinct conformations, and reveal the presence of multiple Packaging Signals potentially involved in assembly regulation that have not been detected by other techniques. XRF also reveals the dramatic effect of the unique Maturation Protein (MP) on both the capsid lattice, and the gRNA conformation inside the phage compared with a virus-like-particle composed only of coat protein subunits. Aspects of genome organisation in the phage, their impacts on the capsid shell, and the distortion of lattice geometry by MP, are hallmarks of molecular frustration. Phage assembly therefore appears to prepare the particle for the next step of the infectious cycle.

## Introduction

RNA molecules play crucial functional roles in cell biology, most of which are dependent on the formation of defined conformational states. The complexity of RNA folding ensembles, however, still challenges our understanding^1^. The relative redundancy of base pairing interactions creates multiple options for the formation of secondary structures. These in turn specify alternatives for formation of tertiary interactions. Many potential conformers are predicted to be of similar folded free energies, implying their potential co-existence. Intermolecular interactions, even with simple cations, can bias folding outcomes^2^, further complicating analysis. It is therefore not surprising that evolution has produced protein chaperones that favour formation of defined folded/functional states of RNAs^3^. Regulated folding is vital in the life-cycles of single-stranded (ss) RNA viruses^4-6^, where genomes are often packaged at a similar density to RNA crystals^7^.

Infectious virions, formed in many cases by spontaneous assembly of viral components, leave one host cell in a state capable of infecting the next. The first step of subsequent infection is often a controlled disassembly of the same viral components. We have shown for a number of ssRNA viral families that their genomes encompass multiple, sequence-degenerate, dispersed sites/motifs termed Packaging Signals (PSs), each of which has affinity for its cognate coat protein (CP) subunit(s)^8-14^. PSs and their interactions with CP subunits collectively ensure sequence-specific, efficient and genetically robust genome packaging^14^. A major implication of this mechanism is that the conformation of the genome (gRNA) adjacent to the protein shell within each infectious particle should be the same or very similar, as predicted by Hamiltonian Path Analysis^15^. For RNA bacteriophage MS2, asymmetric cryo-electron microscopy (cryo-EM)^16-18^ reconstructions have confirmed this suggestion at near atomic resolution (Fig. 1).

**Figure 1.**
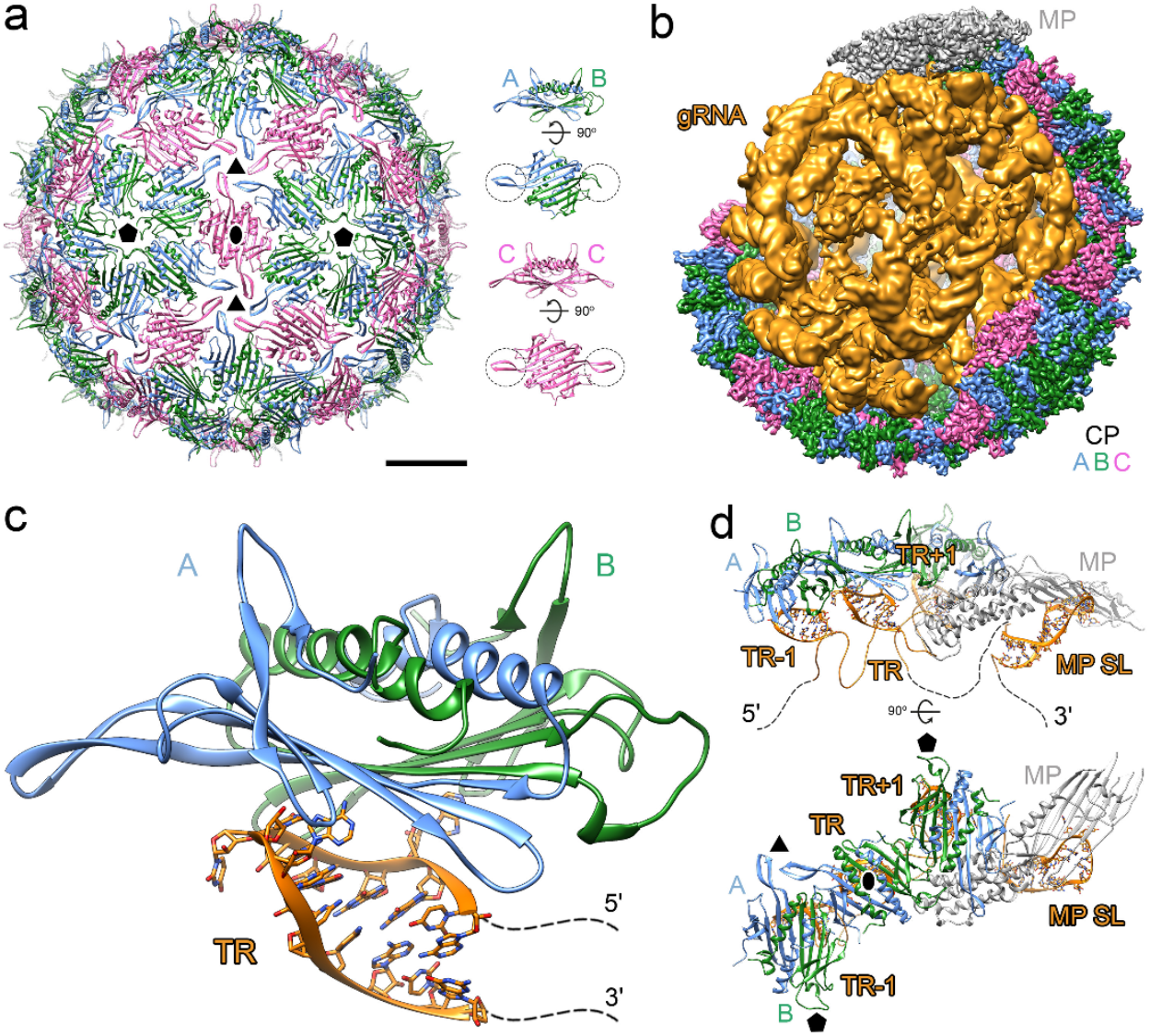
Structure of bacteriophage MS2 & gRNA-protein contacts within the phage particle. a) Atomic model of the front-half capsid of the icosahedrally-averaged crystal structure (PDB: 1ZDH) of MS2 shown with 180 CP subunits as ribbon diagrams coloured in blue (A subunit), green (B subunit) and pink (C subunit) to indicate the differing quasi-conformers, viewed along a two-fold axis^29^. To the right, A/B and C/C dimers are shown, as side views and from the interior of the capsid, with dashed circles highlighting the conformations of the FG-loops. b) Cryo-EM density map obtained without imposing icosahedral symmetry (EMD-8397) reveals the asymmetric organization of MS2^17^, CP colour-coding as in (A), but with one C/C-type dimer replaced by the MP (grey). The internal orange structure, viewed with a section of the capsid computationally removed, is a model of the ordered parts of the gRNA. c) X-ray structure of an A/B CP dimer bound to a nucleotide encompassing TR^52^. Details of the RNA-CP interactions in this structure are shown in Sup. Fig. 1. d) Cartoon image of a portion of capsid seen in (b) (EMD-8397) including the three CP dimers, and their bound RNA stem-loops, and the contacts they make with the MP.

Detecting the formation of sequence-specific gRNA-protein contacts in a large macromolecular machine such as the MS2 phage is challenging. Virion assembly, both *in vitro* and *in vivo* often occurs in seconds^8, 19^, whilst *in vivo* both assembly and genome uncoating often occur within highly complicated cellular environments. We have therefore applied the minimally invasive technique of synchrotron X-ray footprinting (XRF)^20-22^ to determine bacteriophage MS2 gRNA secondary structure and its molecular contacts *in situ*, i.e. within the infectious phage particle. We have also probed how these differ between *in vivo* (phage) and *in vitro* (virus-like particle; VLP) assembled particles. The results reveal that the phage includes multiple examples of molecular frustration, the relief of which likely drive major steps along the infection pathway with a new host.

## Results

### XRF workflow and choice of viral footprinting targets

MS2 bacteriophage (Fig. 1) has a positive-sense, ssRNA genome (gRNA) 3659 nts long^23^. It was the first genome sequenced, and several secondary structures based on enzymatic and chemical footprinting have been determined^24-26^. This gRNA becomes encapsidated in a protein shell (270 Å *dia*.) based on a *T*=3 surface lattice^27-29^. It contains 89 CP dimers (CP_2_) and a single copy Maturation Protein (MP)^17, 30^. The CP dimers exist as two distinct, quasi-equivalent conformers, with 60 asymmetric A/B and 29 symmetric C/C dimers. An additional C/C lattice position is occupied by the MP^30^, the phage component that recognizes the cellular receptor, i.e. the bacterial F pilus^31^. *In vitro* reassembly of gRNA and CP_2_ in the absence of the MP leads to formation of *T*=3 VLPs^32^. This reaction is highly sequence-specific at nanomolar concentrations, a consequence of assembly being regulated by multiple dispersed sites/sequences (Packaging Signals, PSs) across the gRNA^33^.

RNA PSs, which vary in affinity for their cognate CP, define a preferred assembly pathway^13^, biasing the unliganded CP_2_ conformation from C/C to A/B (Fig. 1C)^15, 34-36^. The FG-loops connecting the F & G β-strands in B subunits fold towards the globular body of the protein, in contrast to those in A and C subunits. This allows B subunit loops to fit around the phage five-fold axes (Sup. Fig. 1), facilitated by adoption of a *cis* peptide bond by the essential conserved Pro68 residue^37, 38^. PS encompassing oligonucleotides act as allosteric effectors^33-36, 39^, the replicase translational operator stem-loop, TR^40^, being the highest affinity PS in this gRNA.

XRF is very versatile having unparalleled time-resolution (<100 millisecond exposures) allowing it to analyse RNA folding^21^ and ribosomal assembly processes^22, 41^. It can be used on samples in solution or in the frozen state, and even in living cells. It works by photolysing solvent water molecules, which absorb virtually all the photon energy, creating hydroxyl radicals throughout the sample that differentially modify RNA ribose moieties on the basis of their flexibility (Fig. 2). This leads to cleavage of the phosphodiester backbone at the site of modification making it similar to other footprinting technologies, such as SHAPE^42-44^. Footprinting frozen samples allows direct comparison with the results of cryo-EM. In order to understand the roles of the gRNA in CP_2_ shell assembly, and the impact(s) of the MP on this process, we have determined XRF per nucleotide reactivities of the MS2 gRNA in an infectious virion, in a VLP, and as a protein-free transcript.

**Figure 2.**
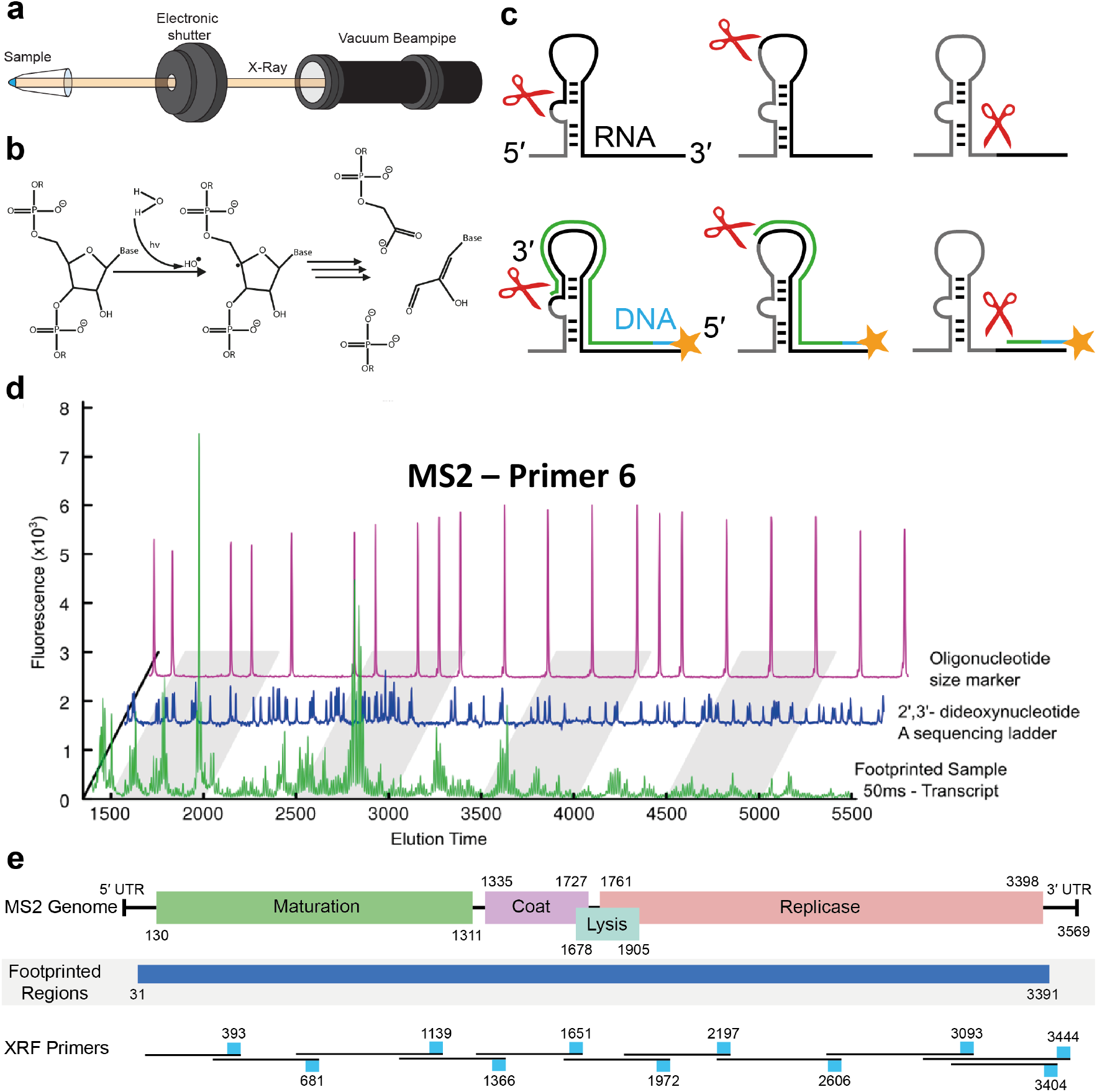
Details of X-ray footprinting across the MS2 gRNA. a) Cartoon showing the arrangement of samples with respect to the X-ray beam. Samples are spun down in Eppendorf tubes, flash frozen and then placed in precisely matched sample holders for exposure. b) The assumed principal chemical reaction leading to polynucleotide chain cleavage. c) A cartoon illustrating the readout of reactivity at each nucleotide by primer extension. d) Example capillary electrophoresis extension traces for a footprinted sample, together with a sequencing ladder and oligonucleotide size markers. e) Genome map of the MS2 gRNA (top) together with the footprinted region (middle) that integrates the primer extensions from the differing primers for both transcript and virion (bottom).

Samples for XRF were flash-frozen, shipped on dry-ice to the Brookhaven National Synchroton Light Source II, Beamline 17-BM, and multiple frozen replicates exposed to X-rays for 10–100 milliseconds. Following exposure, the samples were returned on dry-ice to the host laboratory for further processing. Test exposures and quantitative primer extensions show that avoiding multiple cleavages within the full-length gRNA is difficult to achieve because its different regions cleave at very different rates. We therefore identified conditions that on average yield single cleavages across the region being probed by each primer, assuming that within a frozen sample phosphodiester backbone cleavages outside the transcription product do not cause large-scale conformational changes within the footprinting period.

Eleven reverse transcriptase primers, 19-24 nts long, were identified that provide overlapping transcripts across the region 32-3352 nts for gRNA extracted from exposed samples (Fig. 2e, Table 1). Absorption at 260 nm was used to estimate RNA recoveries and transcript concentrations. Triplicate samples at X-ray exposures of 0, 25, 50 & 100 milliseconds were analysed using a commercial capillary electrophoresis service (DNASeq, University of Dundee) alongside ROX 400HD size-markers (Applied Biosystems™) and reference 2’,3’-dideoxyA sequence ladders (Fig. 2d). Footprinting data only become reliable after an “entry peak” in the electropherograms, creating blank areas adjacent to, and including, the regions where the primers anneal. These genome sequences were analysed using the primer extension reactions from primer sites further 3’ (Table 1). We were unable to obtain extension products with the primer designed to anneal at the 3’ end of the gRNA (3569) or for which extension terminates at the 5’ nucleotide. Both regions are thought to form base-paired stem-loops which may explain these results. The reactivity profiles obtained cover ∼93% of the gRNA, including all but the final 7 nts of the replicase gene, and ∼1000/1048 nts (∼95%) defined as “flexible” in the asymmetric cryo-EM structure due to weak or uninterpretable density^17^.

**Table 1:** XRF Details on Primers - reproducibility and normalization factors. Table shows from left to right: primer IDs (P-IDs), positions, read areas covered by primer extension, average Pearson correlation coefficients (PCCs) over replicates for different states (T transcript, V virion, E extracted and R reassembled/VLP), and normalization factors with standard deviation. The non-normalized per nucleotide reactivities in the region between the first and last size marker of each primer extension were computed as peak height multiplied by peak width for peaks in the footprinted trace; values shown are averaged over replicates.

| P-I | Primer position | Primer extension | Average PCC |  |  |  | Normalisation Factor |  |  |  |
| --- | --- | --- | --- | --- | --- | --- | --- | --- | --- | --- |
|  |  |  | T | V | E | R | T | V | E | R |
| 1 | 393-372 | 31-340 | 0.95±0.018 | 0.86±0.067 | 0.94±0.037 | -- | 1181±20 | 1691±87 | 1886±186 | -- |
| 2 | 681-657 | 279-628 | 0.98±0.003 | 0.90±0.031 | 0.93±0.017 | 0.51±0.204 | 952±23 | 737±38 | 1991±78 | 78±7 |
| 3 | 1139-1119 | 584-1083 | 0.91±0.034 | 0.95±0.006 | -- | -- | 713±107 | 681±65 | -- | -- |
| 4 | 1366-1342 | 966-1315 | 0.94±0.035 | 0.84±0.041 | -- | -- | 378±102 | 979±47 | -- | -- |
| 5 | 1651-1627 | 1251-1600 | 0.90±0.008 | 0.95±0.010 | -- | -- | 99±7 | 1536±130 | -- | -- |
| 6 | 1972-1953 | 1573-1922 | 0.97±0.012 | 0.74 | 0.94±0.030 | 0.57 | 8335±347 | 185±7 | 5347±990 | 108±7 |
| 7 | 2197-2177 | 1796-2145 | 0.80±0.079 | 0.91±0.036 | -- | -- | 808±93 | 218±30 | -- | -- |
| 8 | 2606-2583 | 2137-2556 | 0.83±0.043 | 0.94±0.027 | -- | -- | 130±2 | 124±5 | -- | -- |
| 9 | 3093-3074 | 2542-3041 | 0.94±0.010 | 0.50±0.046 | -- | -- | 2021±93 | 189±8 | -- | -- |
| 10 | 3404-3383 | 3003-3352 | 0.90±0.032 | 0.97±0.009 | 0.28±0.100 | -- | 252±26 | 109±3 | 122±8 | -- |
| 11 | 3444-3426 | 2892-3391 | 0.93±0.023 | 0.51±0.252 | -- | -- | 377±39 | 53±1 | -- | -- |

Open-source software, QuShape^45^, was used to correct the primary electropherograms by applying signal smoothing. as well as baseline, decay and mobility shift corrections. Accurate alignment of peaks and RNA sequence is vital for subsequent analysis. Protection by the virion protein shell typically results in lower signals than with free RNAs, creating problems with sequence matching. We also detected peak alignment issues due to differential mobility of oligonucleotides of different lengths through the capillary. To correct these problems, we developed a series of software algorithms (Sup. Fig 2). All details of data processing are described in Methods.

### Determination of gRNA secondary structures using X-ray reactivities

The resolution of the best MS2 asymmetric cryo-EM reconstruction has allowed direct confirmation of much previous structure prediction in the virion^17^. Experimental verification of secondary structure is essential since its prediction, e.g. by S-fold^46^, on an RNA as long as the genome results in lowest free energy structures that are not meaningful due to the conformational complexity across the ensemble of RNA structures with similar folding free energies. Adding footprinting reactivities or accessibilities as constraints dramatically improves the accuracy of such calculations^47, 48^. We used a well-established approach to incorporate the corrected XRF nucleotide reactivities into secondary structure predictions here, using a scaling factor, *m*, and an offset, *b*, to integrate these into the free energy calculation^43^ (Fig. 3). In previous studies, a single value for these parameters was chosen based on the accuracy of reproducing a known secondary structural element within the footprinted region. The secondary structure for the remainder of the sequence was then calculated based on the same parameter choice.

**Figure 3.**
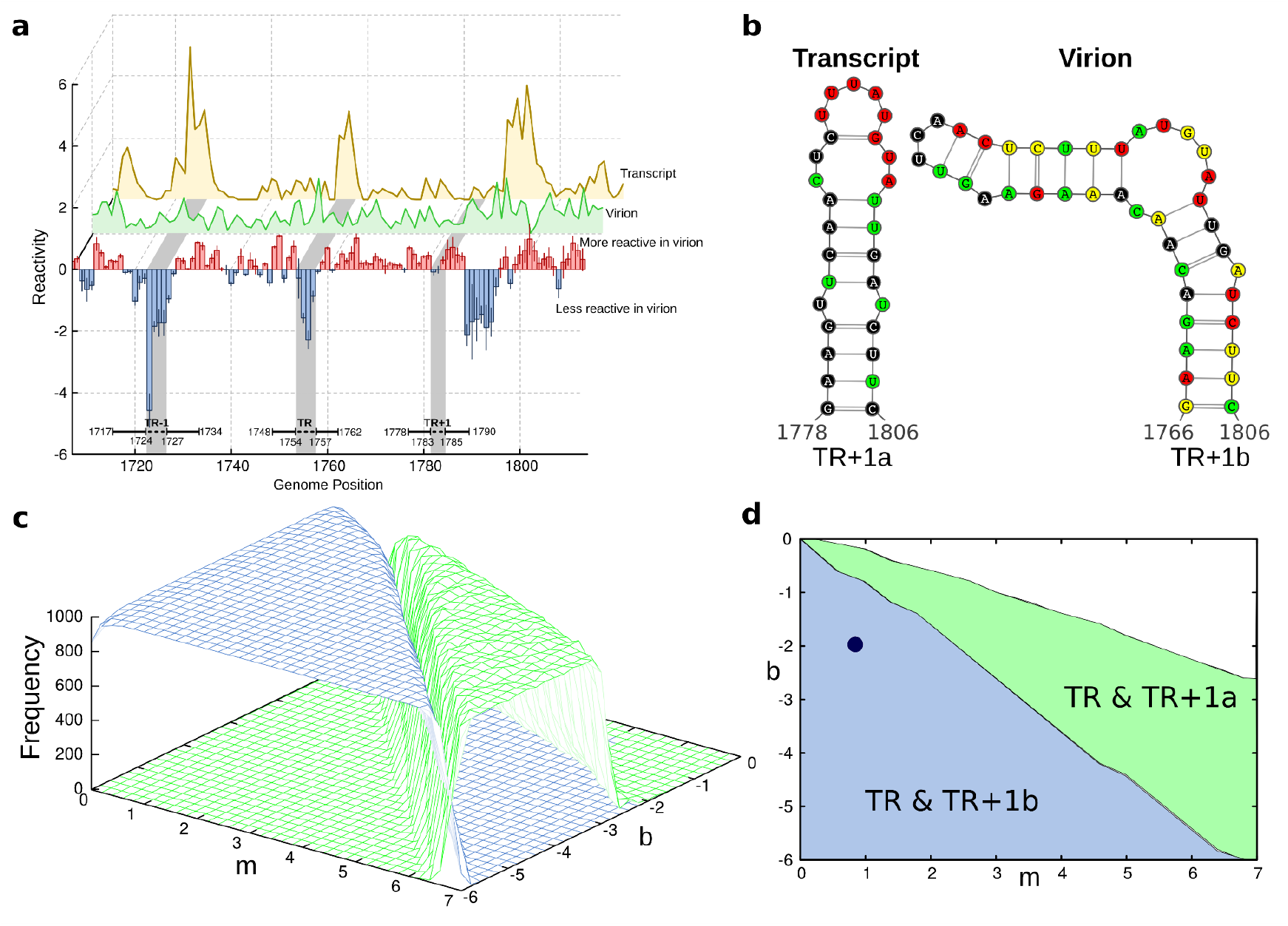
Use of footprinting data to define secondary structures. a) A comparison of the smoothed averages of triplicate XRF reactivities for primer extension reactions with Primer #6 between nucleotides 1700-1820 for transcript (yellow line) and infectious MS2 phage (green line). The relative reactivity differences at each nucleotide between these states of the gRNA are shown as histograms with error bars. Red indicates a site that is more reactive in the virion than in the transcript, whilst blue indictes that the site is less reactive in the virion. b) The calculated S-fold/XRF secondary structures and sequences of the footprinted PS downstream of TR, TR+1. Its preferred conformations in transcript (TR+1a) and virion (TR+1b) are shown confirming the impression from (a) that this sequence undergoes conformational change, creating a PS site in the virion. c) Folding landscape showing the number of times TR and its neighbour TR+1 occur with TR+1 in the TR+1a (blue) or TR+1b (green) conformation in 1000 random structure folds generated for a specific (*m,b*) combination. d) Top view of the folding landscape in (c), partitioning the (*m,b*)-space into areas in which the TR+1a (blue) or the TR+1b (green) is dominant. (*m,b*)=(0.8,-2.0), indicated by a dot, corresponds to the combination at which the number of TR+1a folds (in 1000 random folds) is maximal. This combination is used for secondary structure prediction of the entire genome fold.

The TR stem-loop (position 1748-1762) is readily visible in the asymmetric cryo-EM map^17^ and would be the obvious choice for S-fold parameter selection. To explore this possibility we examined the gRNA footprints (Fig. 3a) from transcript (yellow line) and virion (green line) generated by Primer #6 (Table 1). The lower part of this figure, highlighting the region around TR, shows the differing reactivities of these gRNA states per nucleotide, with error bars for the triplicate determinations. The gRNA is more protected in the virion than in the transcript, as expected. 1000 sample folds across this region, using S-fold constrained by XRF reactivities, at each of the *m* & *b* values shown in Fig. 3c, however, shows that both TR and TR-1 are stem-loops in both virion and transcript gRNAs.These folds are therefore too stable to distinguish virion and transcript folds.

However, the difference map of the raw reactivity data for TR+1 shows different behaviour with a PS-like fold appearing only in the virion. Thus, similar S-fold calculations can be used to identify areas in *m* & *b* parameter space where TR and TR+1a (green), or TR and TR+1b (blue), are the dominant PS-like folds. The *m* & *b* parameters, highlighted as black dots (Fig. 3d & Sup. Fig. 3), yielding these distinct states most frequently across all parameter combinations within the primer read (virion *m* = 0.8, *b* = -2.0, Fig. 3d; transcript *m* = 6.0, *b* = -3.8) were then used for each gRNA for structure prediction (Sup. Fig. 4). The transcript and *in virio* gRNA secondary structures confirm that there are significant conformational differences between them, including folding of PS sites not present in the transcript as predicted^8, 49^. There is minimal variation around a dominant conformation with no evidence for conformational variants, as has been suggested^50^.

**Figure 4.**
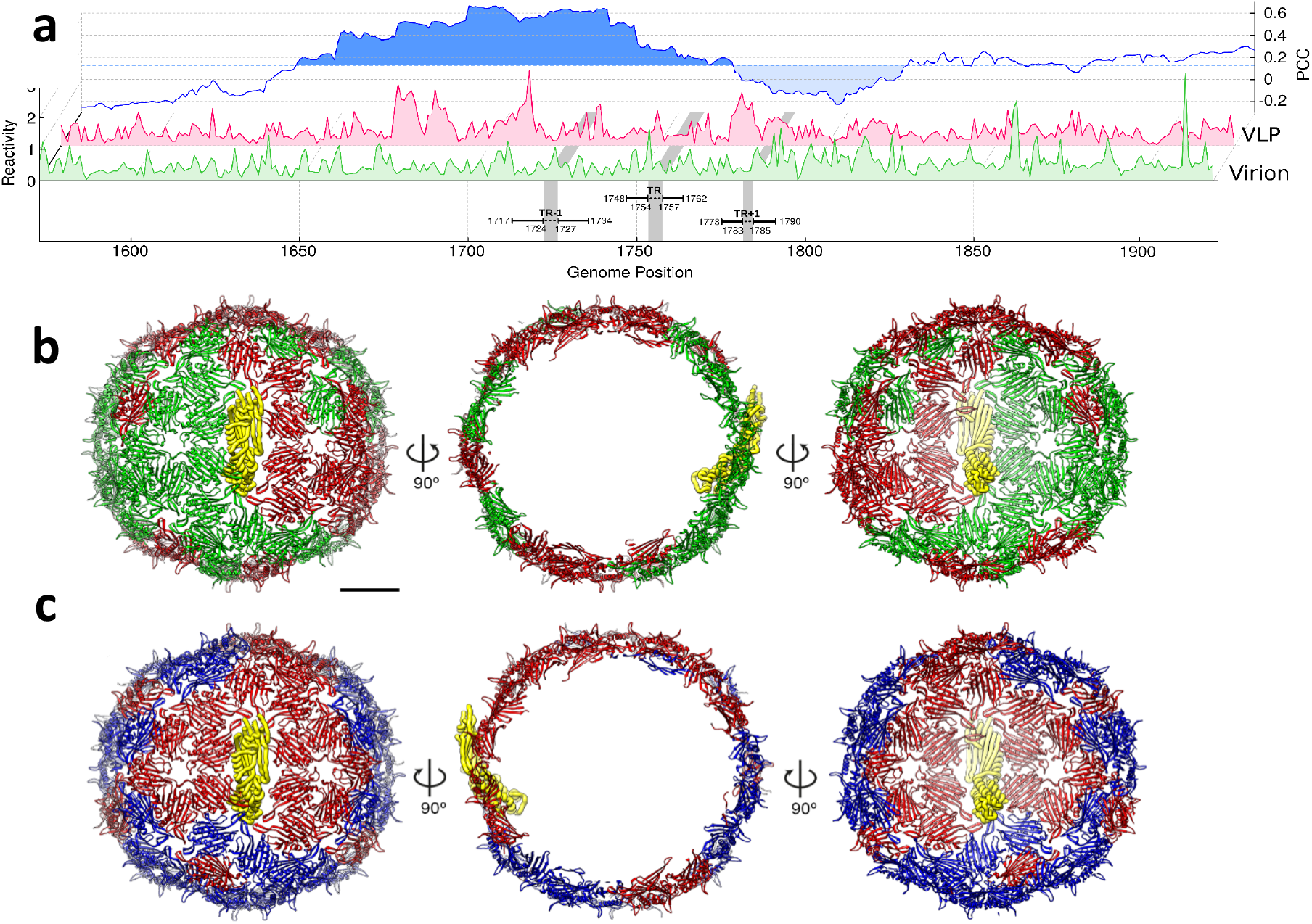
gRNA Secondary structures within infectious MS2 and a transcript. a) PCC comparison of the gRNA *in virio* or in the VLP.(b) Atomic model for MS2 capsid shown as ribbon diagrams with CP_2_-bound PSs (green) and non-bound PSs (red). MP is in yellow. (c) Same as in (b) but with more significantly ‘displaced’ CP_2_ dimers from the idealised *T*=3 lattice (in red) and those less ‘displaced’ CP_2_ (in blue)^17^. The degree of displacement was determined by calculating RMSD between individual dimers in the *T*=3 averaged map and their equivalents in the asymmetric one.

Since RNA phages repurpose their gRNAs from mRNA as assembly substrates, assembly initiation is most likely to occur at the three PS sites centred on TR. In the virion, the MP binds at a specific site adjacent to the 3’ end of the gRNA, and also contacts the CP dimers bound at TR and TR+1^17^ (Fig. 1d). These intermolecular contacts lock the 3’ half (nts 1748-3569) of the gRNA into an RNA loop. There is a well characterised, long-range intramolecular interaction (“Min Jou”) in the gRNA within the virion^23, 51^. For the identification of *m* and *b* values for the transcript from Primer #6 (Fig. 3), which encompasses one half of this interaction, these nucleotides were forced to remain single-stranded, i.e. available for the long-range contact. In the calculation of secondary structures in the virion, this contact emerges naturally, consistent with its appearance in the cryo-EM map.

### gRNA-protein interactions within the virion & infectivity

The XRF reactivity-guided S-fold returns a single secondary structure for the gRNA *in virio* (Fig. 3) with a 97% nucleotide pairing identity across an ensemble of 1000 sample folds, consistent with previous proposals^8, 15^. We assumed that the encapsidated gRNA fold would also reveal the positions of PSs in contact with the CP layer. These need to be distinguished from cleavage protection due to base-pairing which is easy, and RNA-RNA tertiary contacts, which is not.

As a guide for this identification, we looked at the XRF pattern at the strongest natural PS, namely TR at positions 1748-1762^40, 52, 53^ (Figs. 1c and 3, & Sup. Fig. 4). There is no change in secondary structure from the transcript, as expected. There is clear evidence for protection across part of this site, mostly within the tetraloop. The two 3’ nucleotides are more protected in the virion, whilst the nucleotide 5’ to these remains unaffected, and nucleotides both 5’ and 3’ to the loop, and its most 5’ residue, become more reactive. These changes can be rationalised by reference to both solution^54^ and crystallographic structures^52^ for the TR complex (Sup. Fig. 5). Using miminal structure-based assumptions about the 74 stem-loops predicted from XRF-constrained S-fold of the *in virio* gRNA, we identified up to 32 potential PSs having some of the structural features of TR and which are more protected in the phage than in the transcript (Supplementary Material). This value is higher than the 15 tightly-bound stem-loops in contact with the CP shell based on the cryo-EM reconstruction^17^, although those authors point out that a total of 50 stem-loops are in proximity to the capsid and could have acted as PSs during assembly. Note, the XRF criteria set out here exclude several of the PSs identified by cryo-EM. By either method, however, a large fraction of the PSs have dissociated from their CP contacts. The FG-loops in B conformers (Sup. Fig. 1) presumably remain in this conformation in the absence of bound PSs because they are trapped in the protein lattice (Fig. 4), creating local steric preference clashes that will be released upon infection.

Similar events presumably occur in formation of a VLP lacking the MP. However, XRF reveals another fascinating consequence of assembly *in vivo*. The MP imposes a defined conformation on the gRNA between the 3’ end and TR, effectively circularising this part of the gRNA into a loop.. When the Pearson Correlation Coefficients (PCCs) for the XRF reactivities of the gRNAs in phage and VLP are compared (Fig. 4a & Sup. Fig 7), there is reasonable correlation between nucleotides 1650 to ∼1780, i.e. in the region encompassing TR-1 and TR, but this drops dramatically to the 3’ side. Indeed, primer extension data of the VLP gRNA in this region become extremely noisy (Table 1) preventing calculation of a secondary structure. The MP-induced gRNA constrains the interaction between the genome of the protein shell in this region and in its absence it gets scrambled. This result shows that gRNA organisation in a VLP is distinct from that in phage, a differtence revealed for the first time via XRF. Given the importance of RNA-CP contacts for virion assembly, this potentially has significant consequences for VLP assembly and disassembly.

## Discussion

Virion assembly is the critical point of a viral life-cycle, transforming relatively harmless molecular components with great efficiency into a precise molecular machine with the ability to detect, invade and ultimately subvert a host cell. The RNA PS-mediated assembly mechanism^14^ highlights the roles that gRNA-viral structural protein interactions play in this process. These insights focus attention on the conformation of the gRNA within viral particles where, until the advent of asymmetric cryo-EM reconstructions, the presence of ordered RNA fragments was the exception rather than the rule^10, 55^. Even in most of these cases, identifying the nucleotide sequences involved in viral protein contacts is not possible. XRF has allowed us to detect both inter-molecular interactions, such as PS-CP_2_ contacts, and differences in conformations between encapsidated and unencapsidated states of the same genomic RNA with minimal perturbation, i.e. without the need for access of an RNA modification reagent. Its extension to other aspects of viral lifecycles should be straighforward.

XRF of the MS2 gRNA in differing states reveals some of the intimate details of the infectious state with respect to its genome for the first time. To be infectious, assembly of the protective protein shell for a gRNA must be balanced with the need to deliver that genome safely to a new host. This requirement is not usually considered in studies of virion assembly, but some of its molecular details are now directly visible from the MS2 XRF data. XRF reconfirms conclusions based on asymmetric cryo-EM reconstruction^17^, which itself is consistent with PS-mediated assembly^34^, and the inferred Hamiltonian Path^15^ of particle formation. The MP-gRNA complex must form at an early stage of assembly and may play a role in translational repression since the MP blocks access to the 3’ replicase binding site. It also imposes a global distortion across the capsid lattice. The contact to the CP dimer bound at TR will stabilise that interaction but also causes that dimer to sit away from its idealised position within a *T*=3 lattice (Fig. 4c), i.e. like other coat protein dimers adjacent to the MP, it is not in its lowest free energy state.

These insights imply a previously unsuspected aspect of natural virus assembly, i.e. that it simultaneously sets up the conditions of disassembly and gRNA release. The MS2 phage contains multiple structural features that can be described in terms coined by Wolynes and colleagues to describe protein folding^56, 57^, namely “molecular frustration”. They argue that whilst macromolecular systems will globally assume minimally frustrated states (lowest free energy states), locally they must be able to populate highly frustrated states that permit dynamic functions to occur. In the phage context, loss of bound PSs and its impact on the FG-loops in the asymmetric dimers, creates local steric stress in the capsid lattice, the MP imposes an entropic constraint on 3’ gRNA conformation and distorts the icosahedral CP_2_ lattice more globally. All of these contribute to frustration that release of the gRNA-MP complex into a bacterium relieves^31^, i.e. frustration makes infection easier. Such situations more globally in virology would allow virions to “sense” their positions on an infection cycle explaining many aspects of infectivity. It will be interesting to see if the molecular mechanisms visible in a humble phage underlie infections by more problematic pathogens.

## Acknowledgements

We are grateful to Professor Sarah Woodson, Johns Hopkins University, for her encouragement and support in the use of XRF. RT & PGS thank The Wellcome Trust (Joint Investigator Award Nos. 110145 & 110146 to PGS & RT, respectively) for funding, and we also acknowledge the financial support of The Trust of infrastructure and equipment in the Astbury Centre, University of Leeds (089311/Z/09/Z; 090932/Z/09/Z & 106692), and for their additional support, together with The University of Leeds, of the Astbury Biostructure Facility. RT acknowledges additional funding via an EPSRC Established Career Fellowship (EP/R023204/1) and a Royal Society Wolfson Fellowship (RSWF\R1\180009).

Portions of this work used the XFP (17-BM) beamline at NSLS-II. Development of XFP was made possible by the National Science Foundation, Division of Biological Infrastructure (grant No. 1228549), while operations support of XFP was provided by the National Institutes of Health (grant No. P30-EB-009998). NSLS-II, a US Department of Energy (DOE) Office of Science User Facility, is operated for the DOE Office of Science by Brookhaven National Laboratory under Contract No. DE-SC0012704. We thank DNA Sequencing & Services (MRC I PPU, School of Life Sciences, University of Dundee, Scotland, www.dnaseq.co.uk) for DNA sequencing.

## Author Contributions

PGS & RT planned the work, helped to analyse the results and oversaw writing the paper; RC-B, EW, AJPS & AB prepared samples and carried out the footprinting, helping to analyse the results and write the paper with CPM; RJB & SC analysed the data and helped write the paper; EF & JB helped carry out the experiments at Brookhaven.

## Data Availability

The processed data from the capillary electrophoresis data analysis is available as a collection on Figshare (https://doi.org/10.6084/m9.figshare.c.5395302). All other data are available from the corresponding author on reasonable request.

## Code availability

The software package used to analyse the capillary electrophoresis (BoXFP) is freely available to download from GitHub at https://github.com/MathematicalComputationalVirology/XRFanalysis.

## Methods

### X-ray footprinting data analysis

QuShape^45^ has previously been used successfully for the analysis of XRF data of ribosomal RNAs. Due to the significantly longer length of viral genomic RNAs, and additional protection levels arising from their encapsidation inside the viral capsid, this software is not sufficient in our context. We therefore developed a suite of additional algorithms (Sup. Fig. 2), that we implemented in combination with QuShape functions as described below.

#### Initial Data Processing

The CE samples were extracted using the ‘ABIF reader’ tool within the QuShape package^45^, and the following pre-processing steps performed on the raw electropherogram traces:

○ *triangular smoothing* of all traces (footprinted sample, ddA ladder and size markers) for noise removal;
○ *baseline correction* - windows of 60 elution time points were slid in increments of 5 along all traces, and the minimum intensity was substracted from each data point in every frame position;
○ *signal decay correction* for all but the size marker trace;
○ *mobility shift corrections*, mitigating differences in peak positions in the ddA ladder and footprinted sample due to different retention times of the dyes.

These preprocessing steps were performed over 20 windows of the raw electropherograms. The start of the first window was chosen at least 50 elution time points after the entry peak of the electropherogram, and its end point at least 50 elution points before the exit peak, and each subsequent window was obtained via reduction by 5 elution points at either end. The analysis protocol below was carried out for each frame individually, and then the average taken over all frames, in order to mitigate against bias due to the position of the exit peak.

In each case, size marker positioning was performed as an additional preprocessing step. For this, the size marker positions in the electropherogram were determined using the *peak finder* function of QuShape on the size marker trace, and the 21 highest peaks (corresponding to the 21 size markers used) were extracted into the peak list 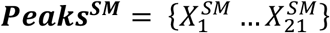.

#### Peak Identification

The *peak finder* function of QuShape was also used to identify peak positions 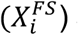 and intensities 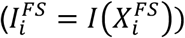 in the footprinted sample trace that were retained in the peak list 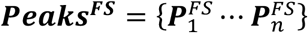, where 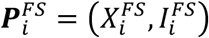. As this method often misses peaks, especially at saddle points and in wide and shallow minima, we developed additional algorithms to mitigate this problem (see Figure S1 I-III):

*Step I:* Saddles were identified as peaks in the negative modulus of the first derivative of the footprinted sample (−|*I*^*FS′*^|). In order to exclude minima, positions corresponding to minima in between peaks were disregarded. Intensity values in the footprinted sample at thus obtained shoulder positions were added to ***Peaks***^***FS***^. In order to identify any missing peaks, the distances between adjacent peaks 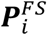 and 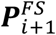 in ***Peaks***^***FS***^, together with their mean (*W*_*m*_) and standard deviation (*W*_*sd*_), were then calculated. For any distances greater than 2*W*_*m*_ − 4*W*_*sd*_, a new peak was inserted at position 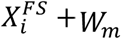 between 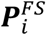 and 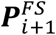 with intensity 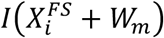.

*Step II*: Any peaks in the footprinted samples (***Peaks***^***FS***^) located between peaks in the size marker peak list (***Peaks***^***SM***^) were exported into a separate list (***Peaks***^***FS***(***sub***)^). The interval between any two adjacent size marker peaks was then partitioned into bins, each representing a nucleotide in the genome, and peaks in ***Peaks***^***FS***(***sub***)^ assigned to these bins. Due to random variation in peak positions along the chromatograms, some peaks will fall into the same bin, and were therefore reassigned to other bins. In particular, for any bins containing no peaks, the closest bin with more than one peak was identified, and their peaks redistributed such that a single peak was assigned to the empty and to the overpopulated bin, each. In case ***Peaks***^***FS***(***sub***)^ exceeded the number of bins, the following procedure was applied: The average over peak positions and intensities associated with each bin was taken; if two bins containing more than one peak were both equally close to an empty bin from either side, then the bin containing a peak position closest to the edge of the empty bin was selected for reassignment into the empty bin.

*Step III:* If ***Peaks***^***FS***(***sub***)^ contains fewer peaks than bins, peaks may be associated with the incorrect bin. This was mitigated via comparison of different replicates. For this, the peak distribution of each replicate was represented as a sequence *B*_*j*_ encoding peak height with reference to the maximal intensity in the ensemble (max (*I*^*FS*(*sub*)^)) as low (L), medium (M), or high (H) as follows:

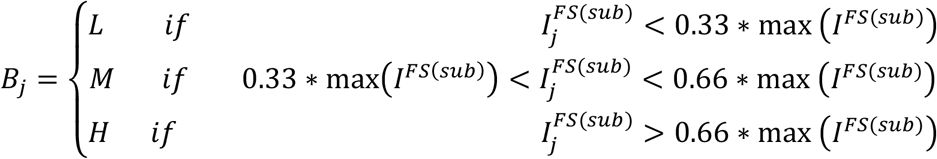

If ***Peaks***^***FS***(***sub***)^ contains fewer peaks than size marker bins, some size marker bins must be identified as unoccupied. This was achieved using a comparison of the sequences *B*_*j*_ of different replicates in the same size marker region, using the Needleman-Wunsch algorithm to create an alignment to identify the positions of any insertions. For this, the following protocol was used:

○ The intensity profiles (*I*^*FS*(*sub*)^) of the peaks in ***Peaks***^***FS***(***sub***)^ were extracted as a sequence;
○ sequences corresponding to different replicates were aligned pairwise using the Needleman-Wunsch algorithm, and a zero value was inserted into the sequences in any position where the algorithm identified an insertion, creating extended intensity profiles;
○ for any two such extended intensity profiles, the Pearson correlation coefficient (PCC) was computed, and for each replicate all its possible extended profiles and PCC value with other replicates retained;
○ the sum over all pairwise PCCs was computed for any possible combination of extended profiles of the replicates;
○ the combination with the greatest sum was chosen as the final alignment, uniquely assigning a specific extended sequence to each replicate; intensity profiles (*I*^*FS*(*sub*)^) were updated accordingly by inserting a peak with zero peak height in the middle of the bin.

The following *final reassignment check* was performed:

○ pairwise PCCs were computed based on ***Peaks***^***FS***(***sub***)^ for the three replicates;
○ if any of these pairwise PCCs was above a threshold, *C*_*PCC*_, the ***Peaks***^***FS***(***sub***)^ of the replicate that did not contribute to this computation was reduced down further by removing bins with an associated peak amplitude of less than 33% of the maximal peak amplitude in ***Peaks***^***FS***(***sub***)^;
○ the above process was repeated for different values of *C*_*PCC*_ from 0.95 down to 0.7 in steps of 0.05;
○ in order to avoid overfitting, alignments resulting in any value migrating more than three bins in ***Peaks***^***FS***(***sub***)^ were discarded;
○ a sequence representation was then generated for each replicate in ***Peaks***^***FS***(***sub***)^ by inserting a peak with zero peak height in the middle of the bin as above.

The individual aligned ***Peaks***^***FS***(***sub***)^ sub lists were then combined into an *aligned peak list for the full primer region* covered by the size marker trace ***Peaks***^***FS***(***part***)^.

#### Reactivity Calculations

Reactivities were calculated for each of the bins in ***Peaks***^***FS***(***part***)^ as the areas under the peaks, 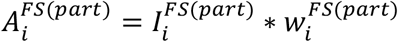, where 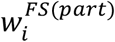 denotes the width of the *i*th peak. Average and standard errors over areas and intensities for different replicate datasets were calculated. In order to achieve a meaningful comparison between reactivies from different footprinted samples, the footprinted peak areas were counter corrected against background using QuShape methods. In particular, *counter corrected reactivities* were obtained as *R* = *Ā* _*tr*_ − *sf* ∗ *Ā* _*b*_, where *Ā*_*tr*_ and *Ā*_*b*_ denote the average peak areas for the footprinted and background samples respectively, and *sf* denotes the scaling factor derived for background samples (0 ms). The standard error was then calculated using trigonometric error propagation as:

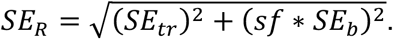

The average over all replicates (*R* values; excluding outliers) and the normalization factor (*nf*) were calculated using QuShape methods. All *R* and *SE*_*R*_ values were then divided by *nf* in order to obtain the normalized area difference *R*^*N*^, and the normalized standard error 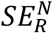.

*Difference plots* were calculated for different states of the gRNA (transcript; *in virio*; extracted; inside VLP) at the optimal exposure time (50 msec, see main text) as follows: The counter corrected reactivities R of both sets were combined, and a normalization factor *nf* determined for this combined set. *R* and *SE*_*R*_ were then renormalized individually using this *nf* from the combined set. The difference values were then calculated as 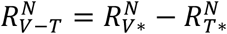, where 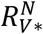 and 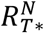 are the renormalized area differences for the individual sets, respectively. Standard errors for the difference maps were again calculated based on trigonometric error propagation applied to renormalized standard error values as above.

We noted that the outcome of the above described procedure may show anomalies if the window used for preprocessing includes the exit peak. In order to mitigate against such errors, we computed pairwise PCCs for the normalised reactivity profiles based on different preprocessing windows. For each window, the average PCC was computed, and any windows with average PCCs of less than 0.8 were discarded. The mean of the normalised reactivity profiles in the remaining windows was used as the final normalized reactivity profile, and the standard error was calculated using trigonometric propagation:

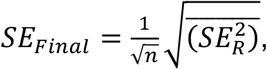

where *n* denotes the number of windows used to calculate the average, and *SE*_*R*_ represent the standard errors of the normalized data for each window.

#### Positioning of the ddA ladder

The positions and intensities of the peaks in the ddA ladder trace were determined using QuShape methods, and filtered into the target peak list ***Peaks***^***SL***^ following the same procedure as outlined for the intensity peaks of the footprinted samples above. In particular, Steps I and II were repeated in order to identify missing peaks in the ddA ladder trace, and for bin reassignment of peaks in ***Peaks***^***SL***^ with positions between pairs of adjacent positions in ***Peaks***^***SM***^. Association of ddA ladder peaks with size marker bins was achieved as follows:

○ If a single peak was located in a bin, the intensity of that peak was assigned to that bin;
○ if more than one peak was assigned to a bin, the average intensity of these peaks was assigned as a single value to this bin;
○ if no peaks were assigned to a bin, then the intensity value of that bin was set to 0.

The individually processed lists (***Peaks***^***SL***(***sub***)^) were then combined into a single aligned peak list for the full primer region (***Peaks***^***SL***(***part***)^), encompassing nucleotides between the start and end of the size marker trace.

In order to correct for errors in peak identification in individual ddA ladders, results were benchmarked against the ladders associated with different replicates (Figure S1 IV). In particular, the intensity profiles of ddA ladders in ***Peaks***^***SL***(***part***)^ were translated into binary sequences *B*_*j*_:

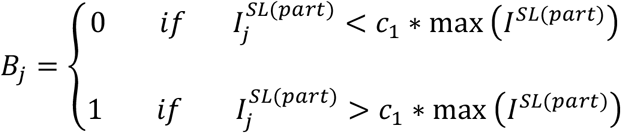

where the constant *c*_1_ was chosen such that the mean PCC over all pairwise combinations of the binary sequences in the ensemble was maximised.

The entries *B*_*j*_ of the vectors ***B*** were then added element-wise into a ‘ballot-box’ array, ***Bbox*** = {*Bb*_1_, ⋯, *Bb*_*n*_}, facilitating comparison between replicate ladders. This array was then converted into a binary consensus sequence with elements 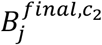:

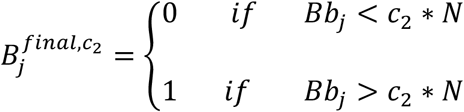

where *N* denotes the total number of datasets in the ensemble, that is, the number of ddA ladders per plate sequenced, and *c*_2_ a constant that we have varied in the range 0.5-0.95 in increments of 0.05. For each fixed value of *c*_2_, this consensus sequence was compared with the genomic sequence in order to determine the optimal position of the ddA ladder. For this, the viral genome sequence was converted into a binary sequence 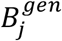, associating a 1 with each occurrence of U and a zero with every other nucleotide:

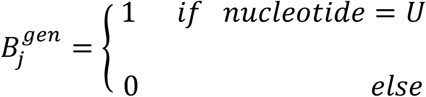

The consensus sequence 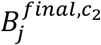 was then slid along 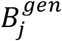 in increments of 1 nucleotide, and for each frame position, the number of correctly matched Us (1s) calculated. For each fixed value of *c*_2_, the maximal number of matches and corresponding frame position(s) were recorded. The position of the ddA ladder with respect to the genomic sequence was then taken to be the frame position with the maximal number of matches across all values of *c*_2_.

## Supplementary Information for

### Additional Biochemical Methods

#### Growth/purchase of MS2 phage

Aliquots (5 μl) of MS2 bacteriophage (ATCC 15597-B1) or *in vitro* transcribed genomic RNA in water were placed in 0.2 ml strip tubes (BrandTech) and flash frozen in liquid nitrogen. The MS2 RNA was transcribed with a T7 kit (NEB) from a pSMART HCAmp plasmid containing a full-length clone (Genbank V00642.1), linearised with *Hpa*I directly 3’ of genome, and subsequently purified by RNeasy kit (Qiagen) as per manufacturer’s instructions. MS2 virion RNA was also extracted from virions using the same kit as per manufacturer’s instructions. Both RNAs were eluted into nuclease-free water (Severn Biotech Ltd) and their integrity confirmed on a denaturing agarose gel (not shown). Sample concentrations were adjusted to a minimum of 200 ng/μl (with respect to RNA concentration in virus samples). All virus and RNA aliquots, including unexposed controls, were stored at -80 °C until shipment for XRF.

#### Determination of XRF Reactivities

The samples were footprinted on beamline 17-BM XFP at Brookhaven National Laboratory^1^, NY, USA. Each beamline session was preceded by a measurement of the relative beam strength. This was achieved via a calibration curve of the photo-bleaching of Alexafluor-488, diluted in 10 mM sodium-phosphate buffer (pH 7.4) exposed to the beam for 10, 15, 20 or 30 ms with 762 μm aluminium attenuation. The fluorescence intensity was measured by fluorimeter for all samples and normalised relative to no exposure sample. The fluorescence was plotted against exposure time and a rate constant, *k*, was calculated. This was used as a measure of relative beam strength which is comparable between runs, and allows sample exposure times to be adjusted to produce similar levels of RNA hydroxyl radical modification between sessions. Samples were mounted in the beamline in a temperature-controlled (−30 °C, ensuring samples remained frozen), 96-well motorised holder, which accommodates individual or strips of 8–12 PCR tubes. Samples were exposed to the beam via a shutter mechanism for 10-100 msec. The holder moves to align each sample tube to the beam for the programmed exposure time. Post-exposure, samples were stored at -80 °C for return shipping to the host laboratory for further processing.

RNA was extracted from exposed virions using Buffer AVL (Viral RNA extraction buffer; Qiagen) for 10 min at room temperature. RNA was bound to magnetic beads (RNAClean XP; Beckman Coulter) and incubated for 10 min. The RNA-bound beads were washed three times with 70% ethanol and allowed to air dry for 5 min, before elution with 12 μl nuclease-free water. The extracted RNAs and beamline-exposed free RNAs were reverse transcribed using Superscript IV (Invitrogen) and a sequence specific 5’ 6-carboxyfluorescein (FAM) labelled primer that attached 3’ of the region of interest. Sequencing ladders were synthesised from in vitro transcribed RNA (2 μg RNA template per ladder) using hexachlorofluorescein (HEX) labelled primers (same sequence as experimental sample primers) and the addition of 1:1 molar concentration of ddATP:dNTP. RNA was degraded with RNase H (NEB, 5 units per sample) and the cDNA purified by precipitation (3x volumes of ethanol, 0.3 M sodium acetate, 0.01x volume of glycogen). Experimental samples were resuspended in 20 μl formamide and sequencing ladders were resuspended in 12 μl H2O, then 1.6 μl of each ladder was spiked into each experimental sample. The samples were heated to 65 °C for 10 min then transferred to a 96-well plate and frozen for shipping to DNASeq (Dundee, UK) for capillary electrophoresis.

### Additional Results

#### Determination of PSs in contact with CPs in virio by analysis of the gRNA XRF data

The strongest PS is TR, and its modification pattern in XRF was used to benchmark other potential PSs. There are differences in reactivity, more or less in the phage, depending on which nucleotide is examined. Sup Fig 1 shows the TR solution structures derived from NMR^2^ and crystallography for the TR in complex with the CP shell (Sup info; Sup Fig 6)^3^. The latter shows that the adenines at positions A1757(A-10) and A1751(A-4) hydrogen bond to amino acids Thr45& Ser47 in A and B subunits, respectively. The designations in brackets are for ease of comparison with previous structural data in which bases were numbered relative to the A(+1) of the AUC replicase start codon. In a TR encompassing oligonucleotide, the A1751 base is known from NMR to intercalate between neighbouring base pairs. The conformational change needed to allow it to make the contacts to a CP subunit could easily destabilise its neighbouring nucleotides increasing their reactivities.

Given the TR-CP_2_ reactivity, we explored every stem-loop present in the calculated encapsidated gRNA structure. There are 74 such stem-loops, 53 of which have an equivalent in the transcript (Sup. Fig. 5). Of these, loop sizes vary from 8(1); 7(2); 6(3); 5(8); 4(27) to 3(12) nucleotides in length. The most frequent, tetraloops, are obviously potential PSs. In TR the 3’ loop nucleotides, U.A, are less reactive in the virion. This is consistent with U1756(U-5) forming a stacking interaction with the side-chain of Tyr85 in the A subunit. We therefore identified all tetraloops (11) in which the two most 3’ loop nucleotides show XRF protection in the virion relative to the transcript, provided there is an equivalent stem-loop in the transcript. This comparison suggests that all 11 are gRNA PSs. There is one example of C.A as the 3’ residues of the stem-loop (UUCA, nts 788 - 791). In isolated oligonucleotides, a cytidine at the 1756(C-5) position makes an additional intramolecular hydrogen bond relative to the wild-type sequence, stabilising the bound conformation and explaining its higher CP affinity^4^. It is therefore not surprising that the XRF reactivity suggests this site is a PS. We then examined the other tetraloops (3) in which a YR (pyrimidine/purine) motif shows low reactivity (i.e. shown as green or black in Sup. Fig.6). For tetraloops without an equivalent in the transcript, we identified those in which the 3’ two nucleotides of the loop (7), or having a YR motif (1), have low reactivity (i.e. on average are smaller or equal to the mean (0.6)).

In total, this identifies 22 of the stem-loops with tetraloops as PSs. An equivalent analysis for triloops and pentaloops, identifies a further 6 PSs for each, making a total of 32 potential PS sites. To confirm these tentative assignments we interrogated the cryo-EM reconstruction rejecting stem-loops involved in RNA-RNA contacts, or a combination of RNA-RNA and CP contacts. This reduced the number of potential PSs sites by 2, leaving 32 PS sites that could have acted during assembly. This list is longer than the 15 sites reported to be tightly bound in the well-resolved regions of the EM reconstruction^5^, and it does not contain all 15 of those sites. Note, XRF reactivities are averages so if there were heterogeneity in the structure, e.g. repeated transient PS binding, the outcome would be misleading. Differences between the lists of potential PSs may also arise because the cryo-EM dataset was repeatedly winnowed to obtain the highest resolution structure. Rather than worry about these interpretations and provisos however, it is clear that multiple PSs within the gRNA remain bound to the CP shell, as expected from PS-mediated assembly^6,7^. It is also clear that many other stem-loop could have acted as PSs and subsequently dissociated from the protein layer.

## Supplementary Tables

**Table Sup 1:** Replicate correlations for transcript. Pairwise Pearson correlations coefficients (PCCs) are shown for correlations of normalized reactivities of replicates A, B and C. The average (rounded to two decimal places) and standard deviations (rounded to three decimal places) over all correlates are as in Table 1.

| Primer | Correlate<br>A/B | Correlate<br>A/C | Correlate<br>B/C | Average ±<br>s.d. |
| --- | --- | --- | --- | --- |
| 1 | 0.947 | 0.936 | 0.972 | 0.95 ±0.018 |
| 2 | 0.983 | 0.978 | 0.977 | 0.98 ±0.003 |
| 3 | 0.906 | 0.946 | 0.878 | 0.91 ±0.034 |
| 4 | 0.919 | 0.923 | 0.981 | 0.94 ±0.035 |
| 5 | 0.897 | 0.899 | 0.911 | 0.90 ±0.008 |
| 6 | 0.983 | 0.962 | 0.962 | 0.97 ±0.012 |
| 7 | 0.758 | 0.893 | 0.753 | 0.80 ±0.079 |
| 8 | 0.877 | 0.792 | 0.822 | 0.83 ±0.043 |
| 9 | 0.946 | 0.933 | 0.952 | 0.94 ±0.010 |
| 10 | 0.865 | 0.928 | 0.908 | 0.90 ±0.032 |
| 11 | 0.957 | 0.924 | 0.914 | 0.93 ±0.023 |

**Table Sup 2:** Replicate correlations for virion. Pairwise PCCs are shown for correlations of normalised reactivities of replicates A, B and C. The average (rounded to two decimal places) and standard deviations (rounded to three decimal places) over all correlates are as in Table 1.

| Primer | Correlate<br>A/B | Correlate<br>A/C | Correlate<br>B/C | Average ±<br>s.d. |
| --- | --- | --- | --- | --- |
| 1 | 0.822 | 0.932 | 0.812 | 0.86 ±0.067 |
| 2 | 0.928 | 0.866 | 0.897 | 0.90 ±0.031 |
| 3 | 0.96 | 0.949 | 0.953 | 0.95 ±0.006 |
| 4 | 0.841 | 0.882 | 0.801 | 0.84 ±0.041 |
| 5 | 0.956 | 0.959 | 0.94 | 0.95 ±0.010 |
| 6 | 0.737 | - | - | 0.74 |
| 7 | 0.886 | 0.953 | 0.897 | 0.91 ±0.036 |
| 8 | 0.932 | 0.926 | 0.975 | 0.94 ±0.027 |
| 9 | 0.475 | 0.466 | 0.55 | 0.50 ±0.046 |
| 10 | 0.963 | 0.976 | 0.958 | 0.97 ±0.009 |
| 11 | 0.32 | 0.414 | 0.795 | 0.51 ±0.252 |

**Table Sup 3:** Replicate correlations for reassembled (R) and extracted (E). Pairwise Pearson correlations coefficients (PCCs) are shown for correlations of normalised reactivities of replicates A, B and C. The average (rounded to two decimal places) and standard deviations (rounded to three decimal places) over all correlates are as in Table 1.

| Primer | Correlate<br>A/B | Correlate<br>A/C | Correlate<br>B/C | Average ±<br>s.d. |
| --- | --- | --- | --- | --- |
| 1/E | 0.978 | 0.909 | 0.922 | 0.94 ±0.037 |
| 2/E | 0.92 | 0.911 | 0.944 | 0.93 ±0.017 |
| 2/R | 0.393 | 0.4 | 0.749 | 0.51 ±0.204 |
| 6/E | 0.975 | 0.928 | 0.918 | 0.94 ±0.030 |
| 6/R | 0.57 | - | - | 0.57 |
| 10/E | 0.254 | 0.19 | 0.386 | 0.28 ±0.100 |

**Table Sup 4:** Correlations between sample conditions. Pairwise Pearson Correlations (PCCs) between the average reactivity profiles for different sample states (T transcript, V virion, E extracted, and R reassembly/VLP) at 50 ms beam exposure for each primer read.

| Primer | T/V | T/E | V/E | V/R | T/R | E/R |
| --- | --- | --- | --- | --- | --- | --- |
| 1 | 0.3 | 0.2 | 0.46 | - | - | - |
| 2 | 0.49 | 0.4 | 0.69 | 0.094 | 0.046 | 0.061 |
| 3 | -0.09 | - | - | - | - | - |
| 4 | 0.07 | - | - | - | - | - |
| 5 | -0.024 | - | - | - | - | - |
| 6 | 0.21 | 0.52 | 0.36 | 0.14 | 0.078 | 0.24 |
| 7 | 0.19 | - | - | - | - | - |
| 8 | 0.009 | - | - | - | - | - |
| 9 | -0.049 | - | - | - | - | - |
| 10 | 0.67 | 0.006 | -0.02 | - | - | - |
| 11 | 0.03 | - | - | - | - | - |

## Supplementary Figures

**Sup. Fig 1:**
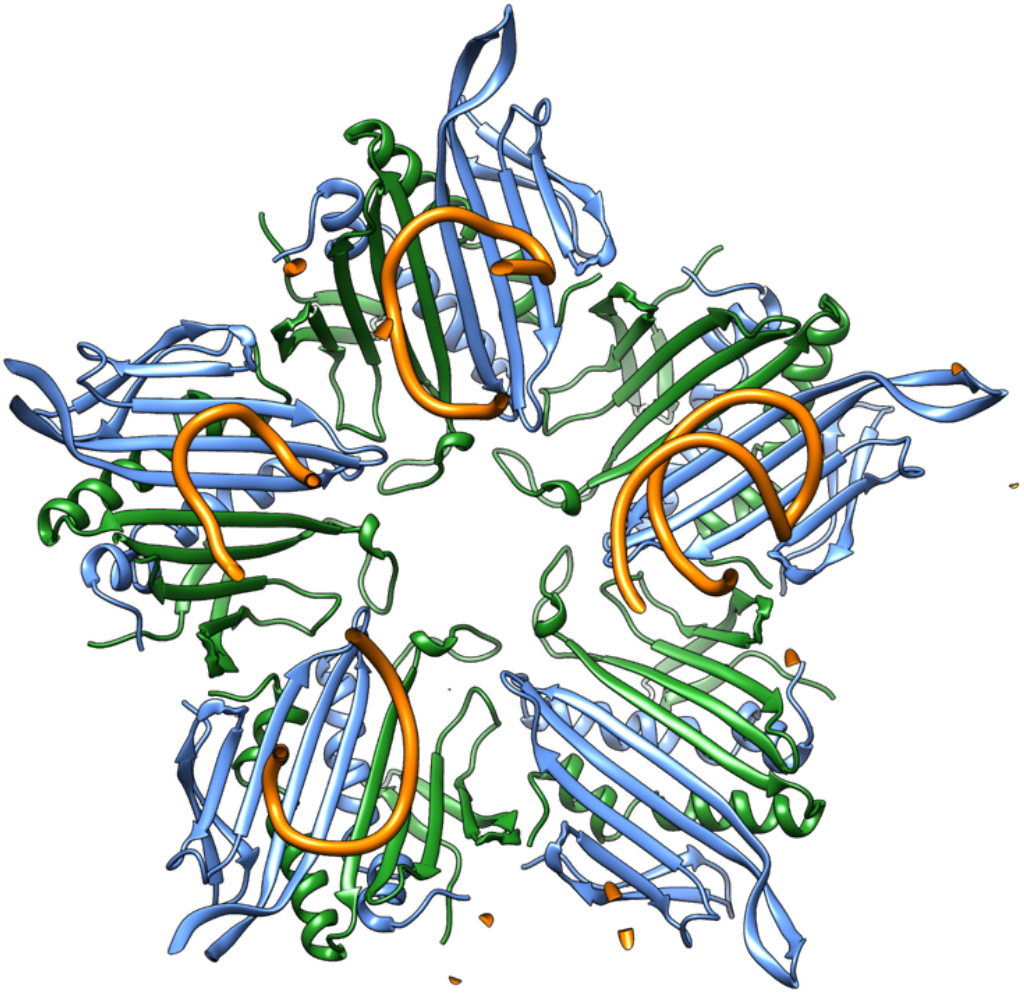
Conformation of FG-loops in A/B dimer lining particle 5-fold axes. View along the 5-fold axis from the phage interior^17^. A/B CP conformers are shown as blue/green ribbons, bound gRNA stem-loops (PSs) are shown as gold rods.

**Sup. Fig 2:**
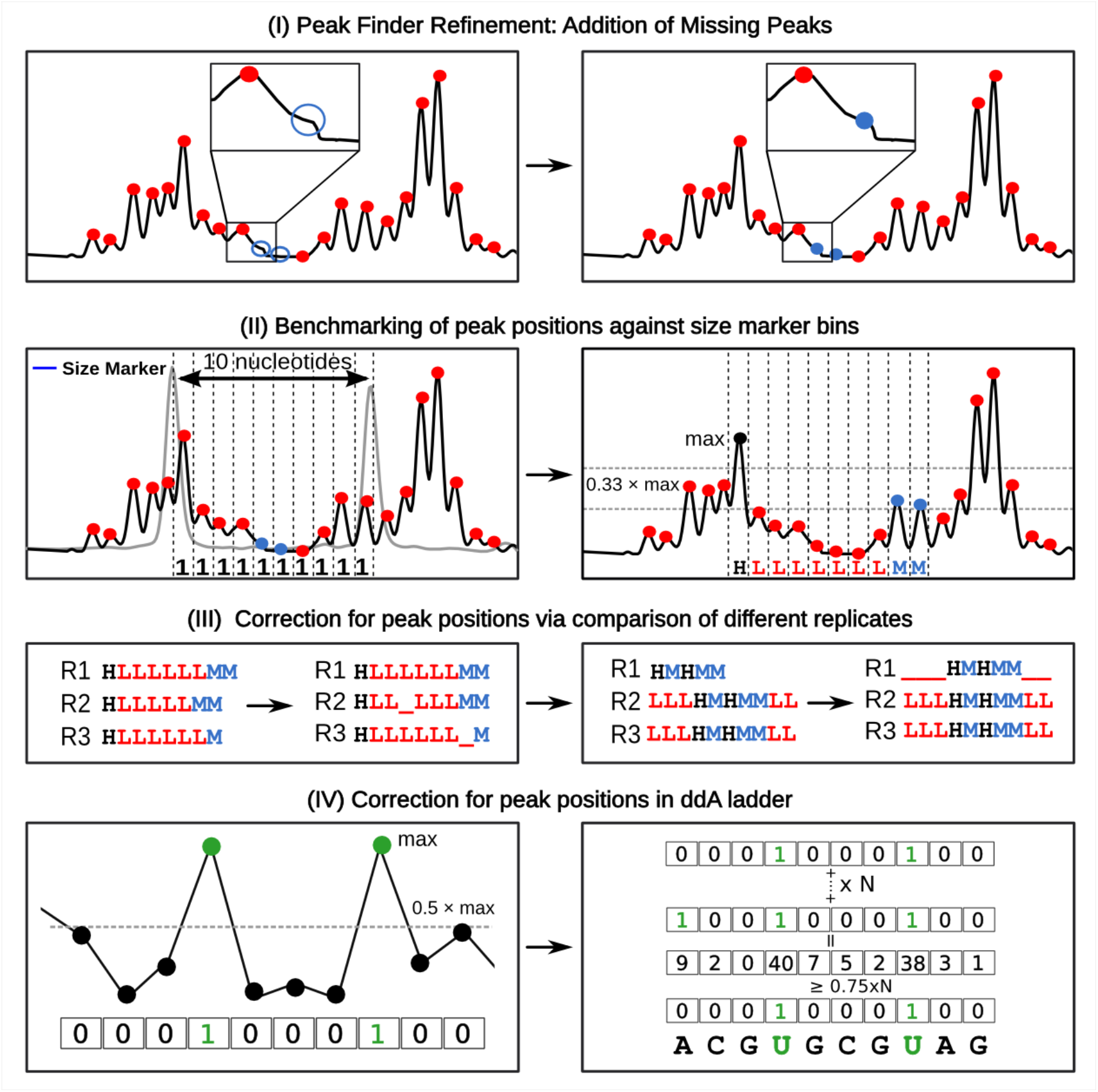
Diagrammatic illustration of peak assigning algorithms.

**Sup. Fig 3:**
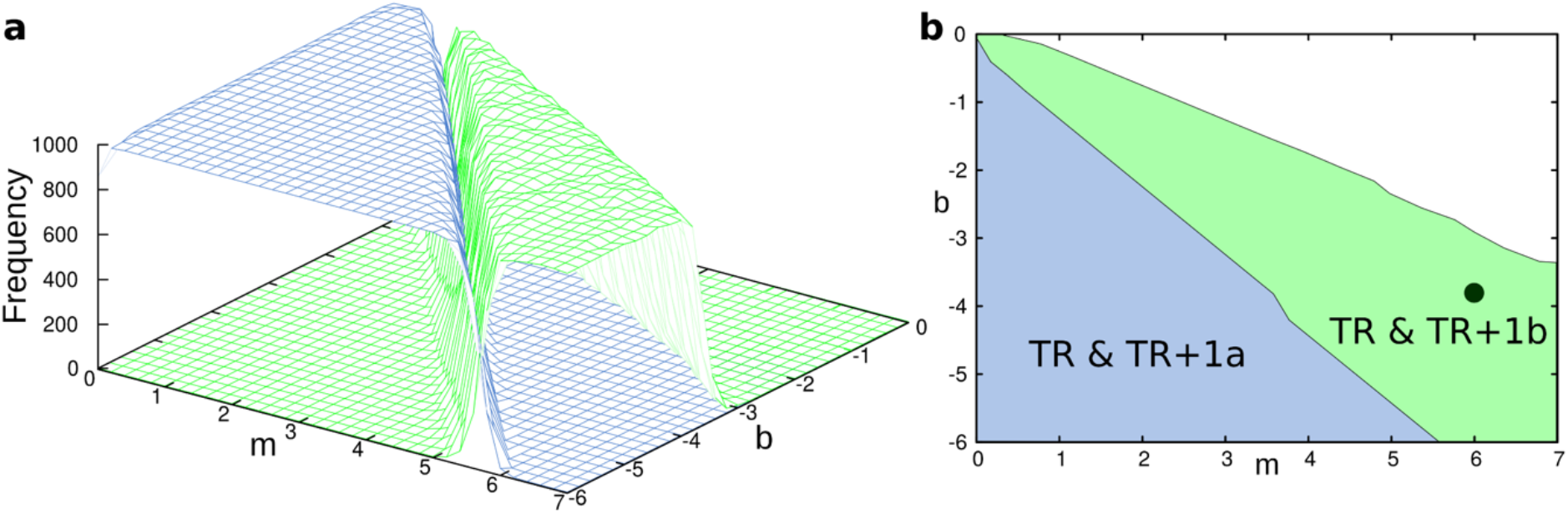
Landscape plot of m & b values for transcript. See also Fig 3 for more details.

**Sup. Fig 4:**
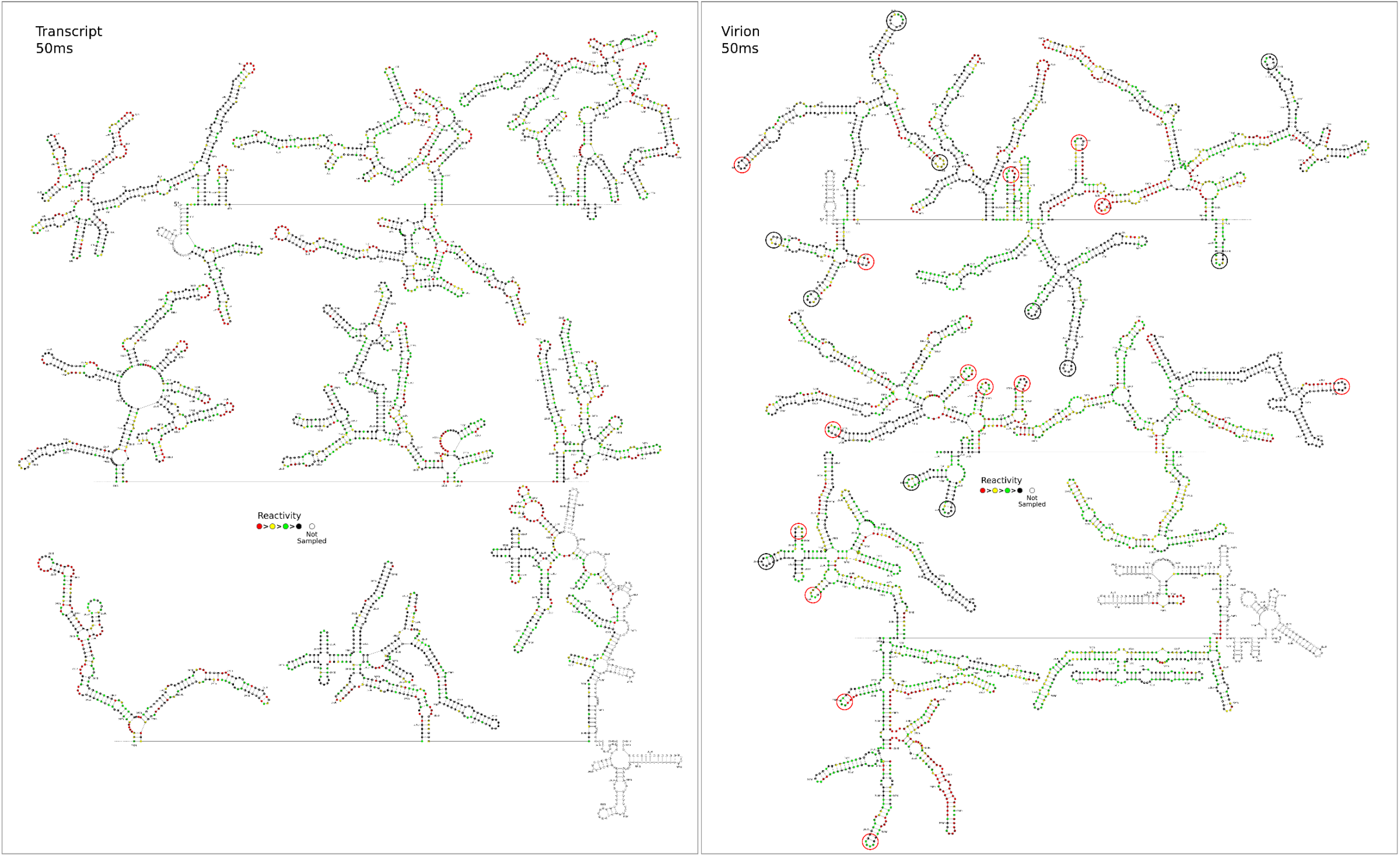
gRNA Secondary structures within transcript & infectious MS2 phage. XRF constrained S-fold structures of the MS2 genome as transcript or *in virio* calculated using the *m* & *b* values from Fig 3 and Fig. 3. Structures are colour-coded to show XRF reactivities, and sites thought to act as PSs are circled in the virion plot. Plots can be enlarged by clicking.

**Sup. Fig 5:**
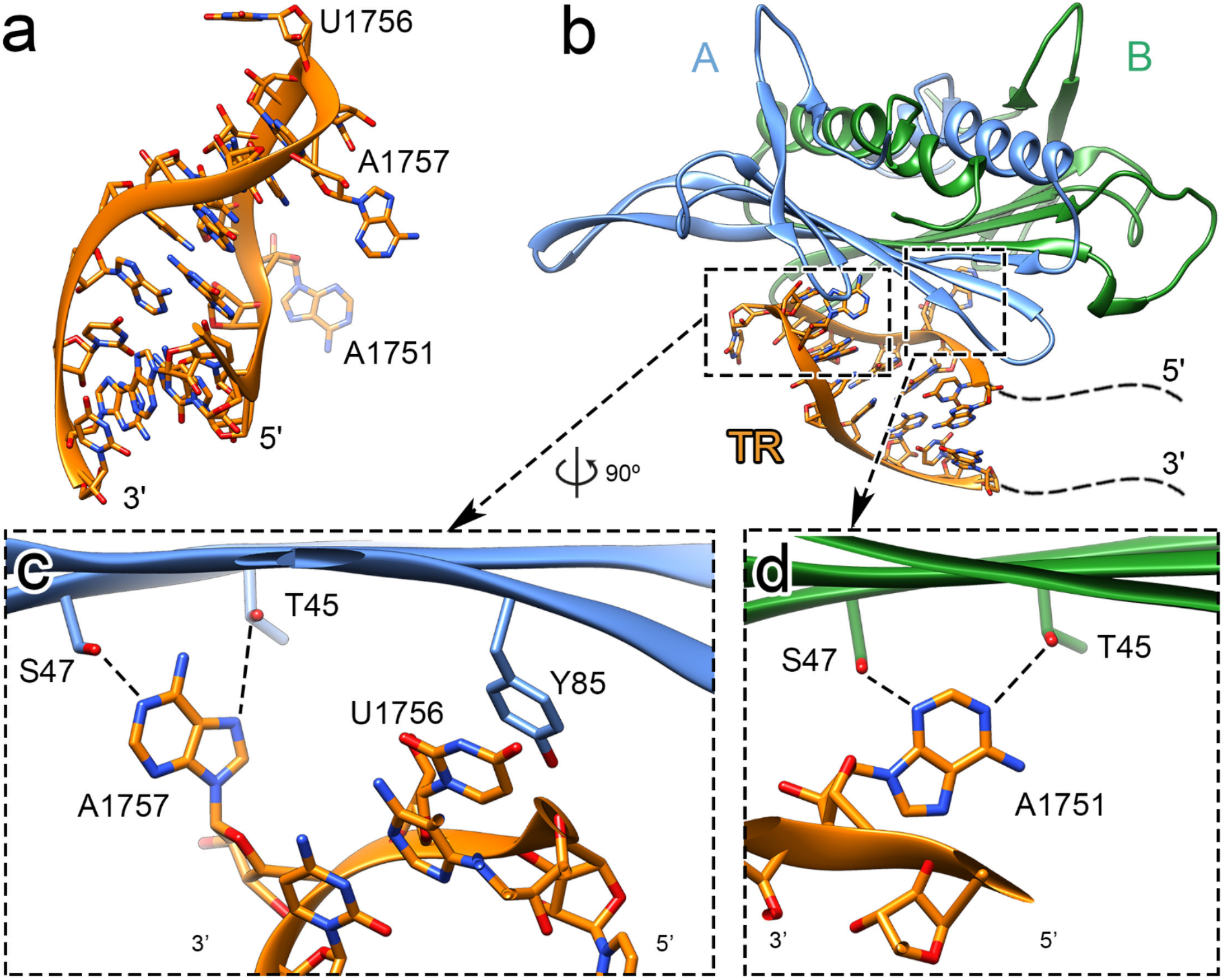
Details of the TR PS and its interactions with an A/B CP dimer. a) The dominant solution conformer of the TR stem-loop seen by NMR^2^. The A-10 base intercalates between neighbouring base-pairs. b) The crystal structure of a TR oligonucleotide in complex with an A/B CP dimer in a VLP. c & d) U1756 (−5) forms a stacking interaction with the A subunit Tyr58, adenines 1751/1757(−10 and -4) hydrogen bond, in two distinct orientations to Thr45 and Ser47 in the two subunits.

**Sup. Fig 6:**
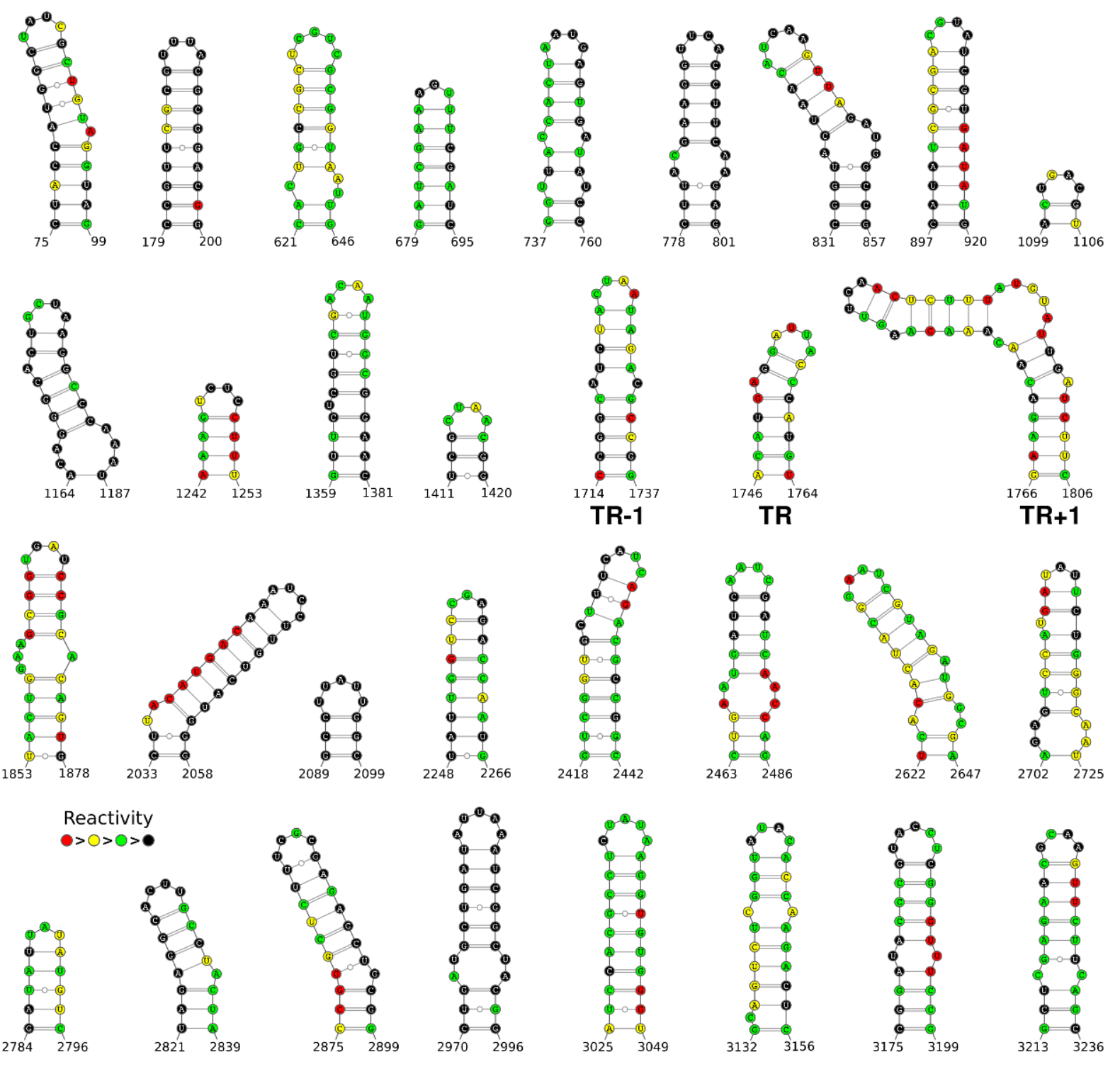
Inferred gRNA stem-loops (PSs) that are bound by the CP shell in virio.

**Sup. Fig 7:**
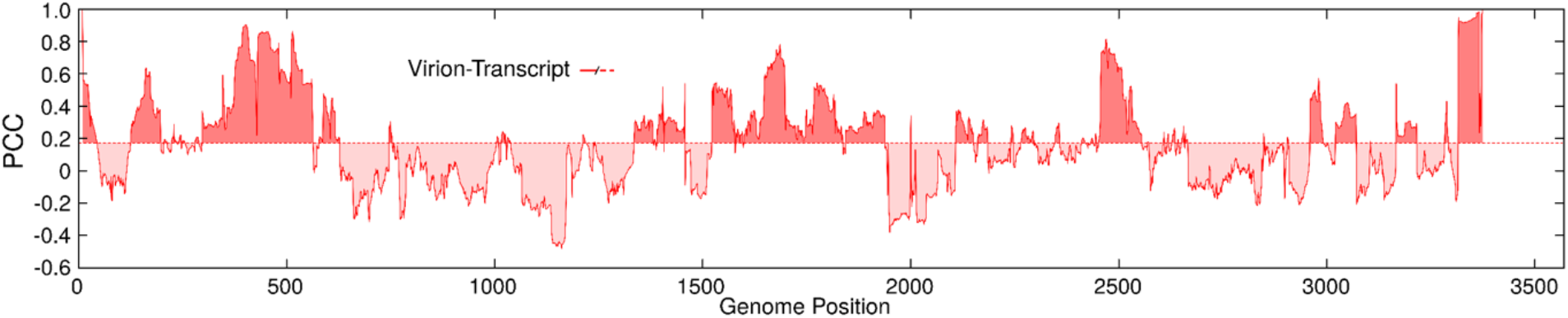
PCC correlation plot. Values for a 50 nt window sliding across the footprinted gRNA region are shown for virion vs transcript, i.e. using data from all the 11 primers (Table 1).

